# Space-Time Light-Sheet Microscopy

**DOI:** 10.64898/2026.04.10.717581

**Authors:** Jinming Zhang, Haokun Luo, Diana M. Mitchell, Shirley Luckhart, Mercedeh Khaja-vikhan, Ayman F. Abouraddy, Demetrios N. Christodoulides, Andreas E. Vasdekis

**Affiliations:** Department of Physics, University of Idaho, 875 Perimeter Drive, Moscow, ID, 83844, USA; Department of Electrical and Computer Engineering, University of Southern California, Los Angeles, CA 90089, USA; Department of Biological Sciences, University of Idaho, 875 Perimeter Drive, Moscow, ID, 83844, USA; Department of Entomology, Plant Pathology and Nematology, University of Idaho, 875 Perimeter Drive, Moscow, ID, 83844, USA; CREOL, The College of Optics & Photonics, University of Central Florida, Orlando, FL 32816, USA; Environmental Molecular Sciences Laboratory, Pacific Northwest National Laboratory, Richland, WA 99354

## Abstract

Light-sheet microscopy (LSM) has revolutionized bioimaging by delivering high-contrast volumetric resolution with minimal photodamage. Spatial wavefront shaping, used to generate lattice and Airy light-sheets, has been particularly effective in advancing LSM beyond the Rayleigh limit. Despite its broad adoption, most LSM implementations rely on rigid dual-objective geometries that complicate sample handling and impose a trade-off between imaging field of view (FoV) and axial resolution. Here, we introduce space-time light-sheet microscopy (ST-LSM), a single-objective strategy that exploits space-time (ST) correlations for the first time. ST-LSM goes beyond separate spatial or temporal modulation to jointly modulate the spatiotemporal spectral structure of a pulse. This uniquely enabled light-sheets with wavelength-scale thickness over millimeter-scale distances. When compared to state-of-the-art approaches, ST-LSM eliminates the dual-objective constraint, expands the sample-accessible volume by 25×, and increases the FoV by 10× without sacrificing sectioning resolution. We demonstrate the versatility of ST-LSM by using a single setup to image specimens across four orders of magnitude in size, from whole roots and developing embryos, down to mammalian cells with sub-cellular axial resolution. These results position ST-LSM as an accessible and high-performance optical microscopy platform at a variety of biological scales, by translating space-time wave-packet physics into a practical imaging modality.

## Introduction

Over the past decades, fluorescence microscopy has permeated most cell biology research because of its exceptional spatiotemporal resolution and contrast.^1, 2^ Among the vast repertoire of techniques, light-sheet microscopy (LSM), also known as selective plane illumination microscopy (SPIM), has transformed bioimaging by delivering rapid, high-contrast optical sectioning of live specimens with minimal photobleaching and phototoxicity.^3–10^ In LSM/SPIM, the illumination objective projects a thin sheet of light orthogonal to the detection axis, restricting the excitation to a single optical plane while preventing unnecessary irradiation of the surrounding regions. This targeted illumination strategy not only significantly reduces photodamage,^11^ but also eliminates out-of-focus fluorescence, thereby enabling the study of dynamic processes in living organisms with enhanced contrast. As a result, LSM/SPIM provides a less invasive alternative to conventional widefield or confocal microscopy, creating new opportunities to observe complex biological phenomena in real time.^12–16^

Despite these compelling advantages, LSM/SPIM faces a critical limitation compared to conventional methods: most configurations require two separate objectives, one for illumination and one for detection. This rigid dual-objective configuration restricts the available space for sample manipulation, severely complicating sample handling and the choice of numerical apertures (NAs) for each objective (Fig. 1a). Consequently, users are often forced to compromise system flexibility and imaging performance. In particular, a high illumination NA is necessary to generate light-sheets that are sufficiently thin (∼1 µm) for subcellular optical sectioning, as commonly reported in related implementations;^5^ however, such objectives typically feature short working distances that restrict optical access to the sample (Fig. 1a).^17^ Moreover, a thin light-sheet is prone to diffraction over short propagation distances,^18^ reducing the usable field-of-view (FoV). Lowering the NA extends the light-sheet length but at the expense of sectioning resolution (Fig. 1b). Balancing configuration simplicity, high sectioning resolution, and large FoV has thus proven challenging, limiting the scope of LSM/SPIM and its combination with widespread sample handling protocols, such as microfluidics.^12–16^

**Figure 1.**
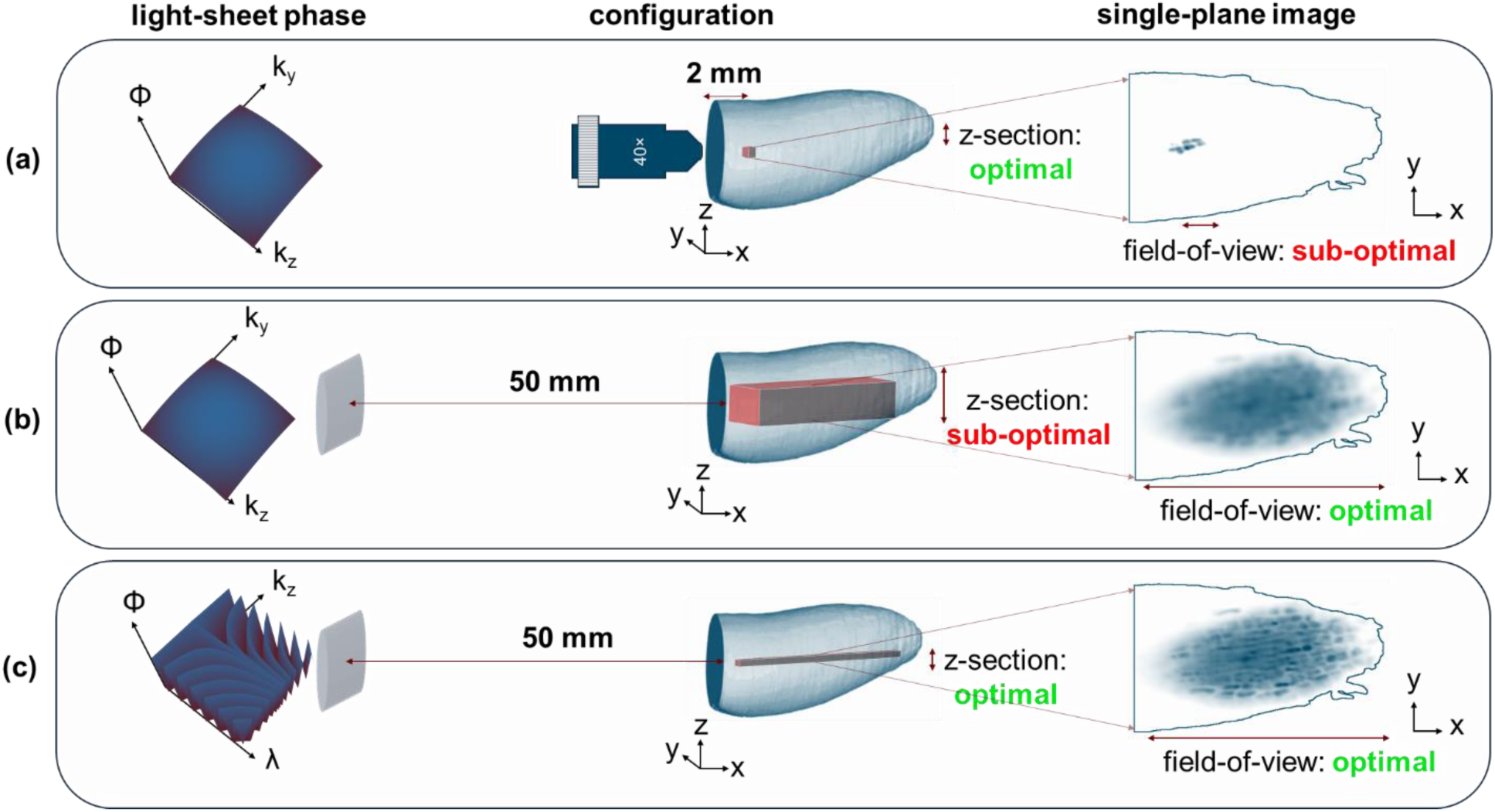
Overcoming the resolution–field-of-view tradeoff in light-sheet micros­copy using ST-LSM. **(a)** Standard Gaussian light-sheet microscopy with a 40× high-NA objective yields optimal axial sectioning but a limited FoV (displayed in the *single-plane image*) due to rapid beam divergence. **(b)** A cylindrical lens extends the illumination prop­agation and enables a wider imaging FoV but degrades axial resolution, resulting in the blurred z-sections shown in the *single-plane image*. **(c)** By integrating spatial (*k*_*z*_) and temporal (*λ*) phase modulation, ST-LSM generates propagation-invariant light-sheets, de­livering both *λ*-scale z-sectioning and extended imaging FoV. Phase-space insets (***Φ***, *k*_*y*_, *k*_*z*_ in **(a)** and **(b)**, and ***Φ***, *k*_*z*_, *λ* in **(c)** illustrate the phase (***Φ***) of the illumination beam struc­ture in each configuration. Root tissue bioimaging examples are included to illustrate the performance in axial resolution (*z-section*) and FoV (*xy-projection*).

To alleviate these constraints, single-objective and open-top variants have been reported.^19–23^ These approaches have enabled considerable progress in relaxing sample handling constraints, yet, they are typically faced with the same axial resolution-FoV tradeoffs. The curved LSM technique has recently enabled impressive FoVs,^24^ offering powerful solutions for large-scale imaging, though it is likely less optimal for applications demanding subcellular axial resolution. Meanwhile, sophisticated wavefront shaping, such as the generation of diffraction-free Bessel, Airy, and lattice light-sheets, generally relying on dual-objective geometries can partially mitigate the axial resolution-FoV tradeoff, yielding remarkable improvements in imaging depth and resolution.^5, 6, 10, 25^

Yet, despite decades of research, achieving 3D subcellular axial resolution together with millimeter-scale FoVs within a single-objective configuration remains challenging, with existing approaches typically optimizing one aspect at the expense of another. Because these beams must remain invariant in both transverse axes, any variant confined in only one dimension to form a sheet inevitably diffracts.^26^ Although it is possible to scan diffraction-free beams to synthesize a light-sheet, the required rapid scanning raises the instantaneous irradiance, which can increase phototoxicity.^11^ This limitation can be overcome by moving beyond purely spatial structuring of a quasi-monochromatic field and instead exploiting the additional degrees of freedom available in broadband pulsed illumination. Finite spectral bandwidth enables a deterministic correlation between temporal and transverse spatial frequencies, thereby permitting propagation-invariant light-sheets under experimentally accessible conditions.^27^ Space-time wave packets (STWPs) realize precisely this type of spatiotemporal spectral correlation, producing propagation-invariant light sheets with symmetric profiles. In addition, STWPs propagate without dispersion in dispersive media,^28^ offer tunable group velocity,^29^ and self-heal after opaque obstacles.^30^ Collectively, these attributes make STWPs strong candidates in LSM, an approach yet to be demonstrated in the context of optical sectioning metrics, illumination-detection geometry constraints, and bioimaging validation.

Building on the unique capabilities of STWPs, we introduce a technique, termed space­time light-sheet microscopy (ST-LSM), that not only resolves the long-standing challenge of dual objective configurations but also overcomes the innate resolution-FoV tradeoff. By leveraging recent advances in spatiotemporal field synthesis,^27^ we constructed a judiciously designed microscope that replaces the conventional high-NA illumination objective with a simple cylindrical lens (Fig. 1c). In this way, we achieve a record 25× increase in the illumination working distance that significantly simplifies the system configuration without sacrificing axial resolution. Importantly, ST-LSM deploys simultaneous temporal and spatial modulation of the illumination beam to produce thin light-sheets (∼1 µm in thickness) spanning millimeter distances, resulting in a substantial expansion compared to standard LSM configurations. This approach opens new possibilities for single-objective light-sheet imaging, where a simple cylindrical lens replaces the high-NA illumination objective, enabling next-generation optical imaging systems with relaxed working-distance constraints.

We rigorously characterized all key aspects of ST-LSM’s performance, confirming its uniform confinement over long propagation distances. We also demonstrated the technique’s considerable versatility by imaging both large-scale structures and subcellular details with the same platform. Specifically, we captured high-resolution, two-photon fluorescence images of intact plant roots and zebrafish embryos, as well as 3D subcellular axial resolution in human red blood cells infected with the malaria parasite Plasmodium falciparum. Collectively, these results demonstrate a purely single-objective strategy that overcomes the sectioning resolution-FoV tradeoff of conventional configurations, paving the way for a new generation of accessible, high-performance light-sheet microscopy platforms. A quantitative comparison of ST-LSM with exemplary Bessel, Airy, lattice, and curved light-sheet implementations, based on reported literature values and evaluated in terms of light-sheet thickness, usable FoV, working distance, and resolution, is provided in the Discussion section.

## Results

### Space-Time Wavepackets

At the core of ST-LSM lies the synthesis of diffraction-free STWPs in the form of light-sheets, an emerging class of structured optical pulses that exhibits both propagation invariance and dispersion immunity.^27, 28^ Unlike conventional beam shaping techniques for generating optical lattices, Bessel, and Airy beams,^31–33^ or separable spatiotemporal modulations, such as the generation of Airy-Bessel light bullets,^34^ STWPs are synthesized by introducing a prescribed spatiotemporal spectral structure into a pulsed optical field, whereby each temporal frequency (or wavelength) is tightly associated with a distinct transverse spatial wavenumber.^27^ This coupling fundamentally departs from purely spatial or separable spatiotemporal strategies, with benefits previously demonstrated in extended propagation distances;^27^ here, we extend these advantages to biological imaging and in the context of LSM/SPIM for the first time.

By contrast, conventional LSM/SPIM configurations (Fig. 1a) typically deploy a Gaussian light-sheet with a small beam waist (w_0_) to enhance optical sectioning. This, however, comes at the cost of a reduced FoV, with the Rayleigh length scaling as x_R_ = π · w_0_^2^/λ_0_ (λ_0_ is the central wavelength of the pulsed light-sheet). Further, a narrow w_0_ demands a high-NA focusing lens,^17^ which limits optical access to the sample due to its short working distance (Fig. 1a). Conversely, increasing the beam waist w_0_ (i.e., lowering the illumination NA) enlarges the FoV but sacrifices axial resolution and optical sectioning efficiency (Fig. 1b). As such, purely spatial approaches must navigate the innate trade-off between tight axial confinement (z-axis in Fig. 1b) and extended coverage (x-axis in Fig. 1b).

To overcome this trade-off, ST-LSM harnesses the exotic properties of STWPs, which suppress diffraction while maintaining high axial confinement. STWPs achieve this through the deterministic correspondence between the spatial and temporal frequencies introduced into the structured light field.^27^ Specifically, the ST light-sheet employed in this work can be described by a spectral constraint of the form 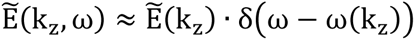, where 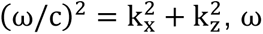 denotes the temporal frequency, and k_x_ and k_z_ represent the longitudinal and transverse wavenumbers, respectively. Note that we use x here for the axial coordinate of the STWP light sheet and z for the transverse coordinate, as appropriate for LSM, in contrast to the previous literature on STWPs where x and z are exchanged. For convenience, the dispersion relation ω = ω(k_z_) is defined by the intersection of a plane and the surface of the free-space light cone 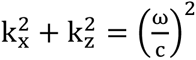, as displayed in Fig. 2; here θ is the angle the plane makes with the k_x_-axis, which is referred to as the spectral tilt angle. Fig. 2a depicts the broad “patch” characteristic of a conventional Gaussian pulsed beam, highlighting its susceptibility to diffraction and dispersion through the k_x_ dependence on k_z_, as displayed in the projection onto the (k_x_, k_z_)-plane.

**Figure 2.**
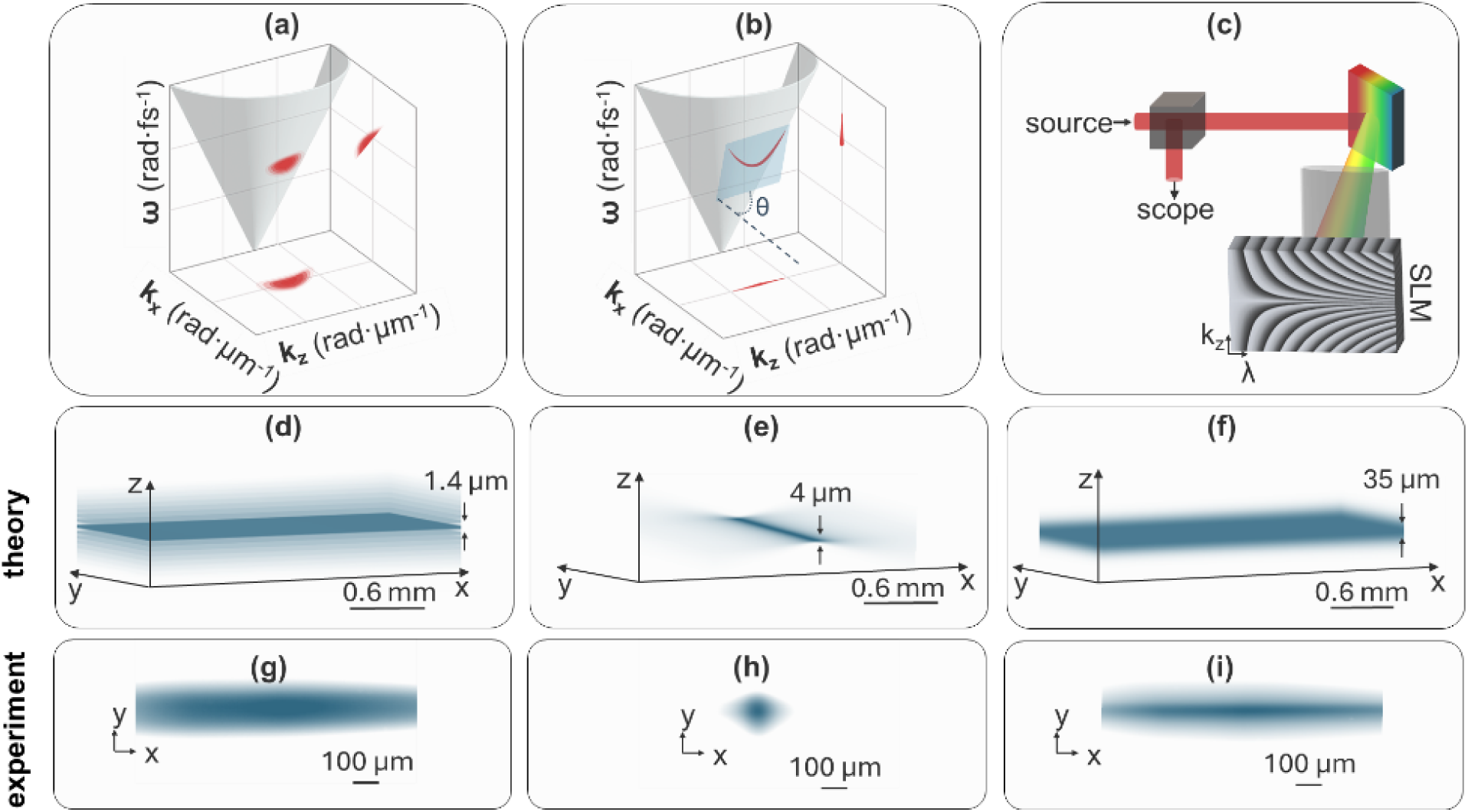
Space-time (ST) beam shaping enables propagation-invariant, ultrathin light sheets. (a) A pulsed Gaussian beam represented as a broadband spectral patch on the *k*_*x*_, *k*_*z*_,and *λ*light-cone that leads to diffraction in real space. (b) A hyperbolic ST spec­tral trajectory correlating transverse spatial frequency (*k*_*z*_) and wavelength (*λ*), yielding invariance of *k*_*z*_with respect to the propagation constant *k*_*x*_; *θ*denotes the spectral tilt an­gle defining this conic-section trajectory. (c) Schematic of the pulse shaper used to syn­thesize ST light-sheets via a 2D phase mask encoding the *k*_*z*_–*λ* correlation on the SLM. (d-f) Computed *yz*-plane propagation of (d) an ST light-sheet (∼1.4 μm FWHM) sustained over ∼1 mm, (e) a Gaussian light-sheet (∼4 μm waist) exhibiting rapid divergence, and (f) a cylindrically focused beam extending propagation at the cost of axial confinement (∼35 μm). (g–i) Corresponding experimental transverse (*xy*) intensity profiles at focus for the three cases.

Building on this framework,^27^ we synthesized tightly focused light-sheets with near-wavelength axial thickness that tightly focus through a simple cylindrical lens, rather than an objective, and maintain their profile over millimeter-scale distances. In the paraxial, narrowband regime, angular dispersion control is utilized to associate each temporal frequency ω undergirding the optical pulse profile with a transverse wave number (or spatial frequency) k_z_ undergirding the transverse beam profile (Methods), governed by the rela­tion:

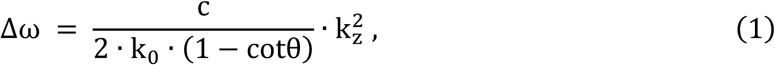

where the bandwidth Δω is Δω = ω − ω_0_. In this context, the selected tilted (hyperbolic for θ > 45°) dispersion trajectory (Eq. 1) projects as a narrow line in the (k_x_, k_z_) plane (Fig. 2b) in stark contrast to the broadened patches of Fig. 2a associated with a standard Gaussian pulsed beam.^23^ This stringent correlation is the key to diffraction-free propagation, introducing millimeter-scale propagation distances in ST-LSM, while maintaining near-wavelength-scale optical sectioning.^35^

### Single-Objective Light-Sheet Microscopy

To generate the STWP, we first dispersed a femtosecond laser pulse with a diffraction grating (Fig. 2c), then directed the spatially resolved spectrum, via a cylindrical lens onto a reflective, phase only spatial light modulator (SLM). This pulse shaping configuration, further detailed in Fig. S1 and comprising multiple auxiliary optical elements required to efficiently generate an STWP required to efficiently generate an STWP, enables precise encoding of one-to-one mapping between each temporal frequency ω and its corresponding transverse spatial frequency k_z_. The retro-reflected wavefront retraces a folded path through the same cylindrical lens back to the grating, whereupon the pulse is re-constructed, thereby, preserving the generated spatiotemporal structure.^36^ Two pairs of cylindrical lenses then relayed the structured STWP onto the imaging plane: the first pair (CL_y1-2_, Fig. S1) compressed the beam along the y-axis (Fig. 1c), and the second (CL_z1-2_, Fig. S1) along the z-axis (Fig. 1c). Once formed, a cylindrical lens (CL_z2_ in Fig. S1) delivered the ST light-sheet into the specimen with a 50 mm working distance. This distance represents more than 25× increase over standard high-NA illumination objectives, greatly expanding the possibilities for sample handling, including more flexible mounting geometries.

To collect the corresponding light-sheet fluorescence image, we implemented a standard SPIM configuration (Fig. S1) using a single water-immersion detection lens.^9^ We characterized the light-sheet emerging from CL_z2_ at the sample plane by directly imaging its intensity along the yz-plane (facet imaging, Fig. 1c) and by capturing its propagation characteristics via two-photon fluorescence in an appropriate medium (Methods) along the xy-plane (Fig. 1c). As shown in Fig. 2d-f, these results fully corroborated the theoretical predictions, demonstrating both the propagation-invariant behavior of the ST light-sheet along the x-axis and its near-wavelength-scale thickness along the z-axis (Fig. 1c). Specifically, we measured a ∼1.4 µm full-width-at-half-maximum (FWHM) thickness, sustained over a propagation distance of more than 1.2 mm ± 35 μm (Fig. 2d and Fig. S2a). In essence, this corresponded to an approximately 0.15 mm^2^ effective FoV at 16.7× imaging magnification and covered the complete image frame at 40× magnification. By contrast, a standard Gaussian beam delivered through a high-NA illumination objective propagates only 147 ± 1 μm (Fig. 2e and Fig. S2b) with markedly lower optical sectioning because of the beam’s 4 μm wide waist along the z-axis (Fig. 1a). Concomitantly, a low-NA illumination lens can achieve a comparable propagation distance but at the cost of broadening the sheet waist to ∼35 µm (Fig. 2f and Fig. S2c), highlighting the unique advantage of our ST approach.

To determine the effective axial resolution, we measured the 3D point-spread function (PSF) using 0.2 µm fluorescent microspheres embedded in agarose (Methods). Volumetric stacks were acquired with the 40×/0.8 water-immersion detection objective, and axial intensity profiles were extracted from isolated beads. Gaussian fitting yielded an average axial FWHM of 1.4 ± 0.1 µm (n = 9 beads) (Fig. S2d). Because the bead diameter is substantially smaller than the measured axial width, convolution-induced broadening is limited (<10%), indicating that the measured PSF closely reflects the optical sectioning performance of ST-LSM. To assess the spatial uniformity across the FoV, we quantified bead-based axial FWHM at multiple lateral positions spanning the light-sheet, including near-edge regions (Fig. S2e). The extracted values indicate preserved axial confinement across the full lateral extent of the sheet. Importantly, the extended propagation of the ST light-sheet does not imply that the optical power is distributed over a longer axial distance. Instead, because the ST beam suppresses diffraction, the local peak intensity along the propagation axis remains comparable to that of a Gaussian light-sheet of comparable waist and pulse energy.

All propagation and PSF characterizations above were performed in homogeneous media to isolate the intrinsic diffraction properties of the ST light-sheet. In scattering biological tissue, attenuation due to scattering and absorption ultimately limits imaging depth, as in all optical microscopy modalities. Importantly, prior work has shown that STWPs exhibit enhanced resilience to scattering due to their underlying space-time spectral correlations (“classical entanglement”).^35^ We independently verified this behavior experimentally (Fig. S3), demonstrating that the ST light-sheet exhibits an increased effective propagation length in scattering media relative to a Gaussian reference under comparable conditions.

As recently reported,^27^ STWPs often produce substantial side lobes, namely local intensity maxima surrounding the main lobe due to interference among the STWP frequency components. This effect is reminiscent of Bessel and Airy beams,^5, 6^ whose diffraction-free propagation partly arises from their inherent side-lobe structures. By combining modeling and experiments (Methods), we found that different STWP trajectories on the light cone (Fig. 2b) alter both the main-lobe width and side-lobe intensity (Fig. 3). These two metrics are inversely related, so that narrowing the main lobe increases the intensity proportion of side lobes (Fig. 3). The latter stems from the Fourier relation between the frequency domain and pertinent spatiotemporal profiles.

**Figure 3.**
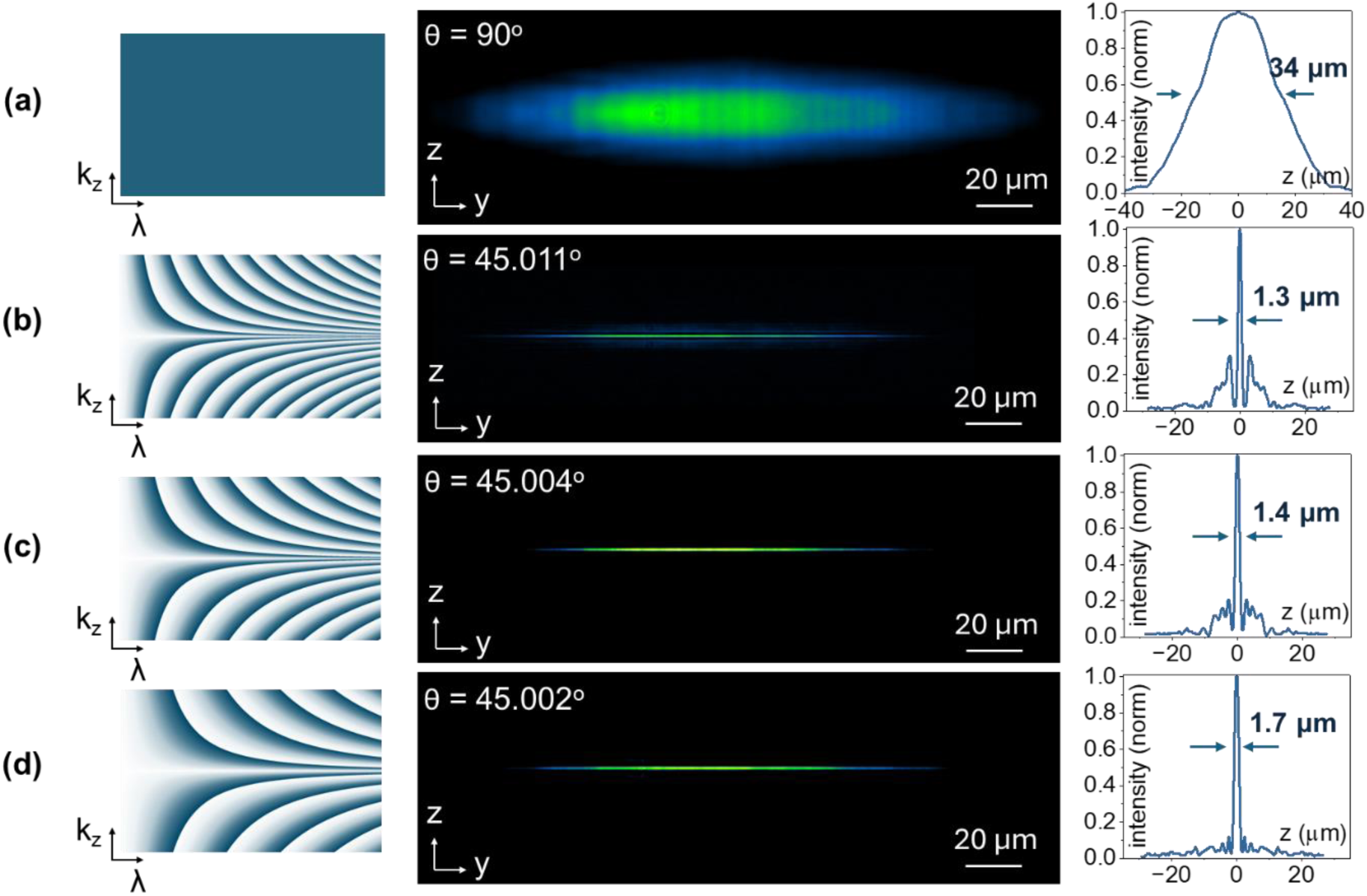
Light-sheet synthesis via space-time (ST) phase modulation. **(a)** Conven­tional Gaussian beam with no phase modulation yielding a 35 μm-thick light-sheet (*right*), as shown in the measured *z–y projection* (*center column*). **(b)** A STWP with a moderate phase gradient generates a thinner beam (∼1.2 μm), though with considerable side-lobe intensity. **(c-d)** Decreasing the angular spread of the STWP modulation further sharpens the beam, yielding axial thicknesses of ∼1.4 μm and ∼1.6 μm, respectively, however, with enhanced side-lobe suppression. *Left column:* depicts the phase masks used to synthe­size each ST with the respective θ values displayed in the intensity profiles (*middle col­umn)*; *right column* plots the extracted intensity cross-sections across the *z-axis* at the beam focus, confirming the wavelength-scale optical sectioning in ST-LSM configurations.

Through this analysis, we identified an optimal configuration for two-photon light-sheet imaging at ∼1.4 µm (FWHM), where the side lobes remain below ∼10% of the main-lobe’s peak intensity (Fig. 3c). Notably, further reductions to the side-lobe intensity forced the main lobe to broaden, thereby weakening the optical sectioning and overall imaging efficiency by introducing excessive out-of-focus excitation (Fig. 3d ). This 1.4 µm – 10% balance demonstrates how fine control over ST trajectories can achieve powerful imaging performance without requiring deconvolution,^6^ a key advantage in bioimaging applications, as discussed in the following section.

#### Bioimaging

To showcase the versatility of ST-LSM in bioimaging, we first examined live roots from the model legume Medicago truncatula, a challenging specimen due to its dense cell walls and millimeter-scale size. The enhanced FoV of ST-LSM allowed us to visualize in a two-photon format substantial segments of root tissue in a single acquisition, revealing both overall architecture and individual cells within the epidermal (“skin”) layer. By employing 3 µm step sizes along the z-axis (Fig. 1c), we could image millimeter-long root sections in under 30 minutes when using 1 second integration times per frame. This step size was selected to balance volumetric coverage and acquisition speed and does not correspond to Nyquist sampling of the measured axial PSF. Importantly, this exposure time was chosen conservatively and does not represent a fundamental limitation of ST-LSM. The total acquisition time was dictated primarily by stitched field size and sampling density rather than beam formation constraints, consistent with volumetric imaging strategies employed in other two-photon light-sheet platforms.^10, 37^ Shorter exposure times can be implemented depending on fluorophore brightness and detection conditions, enabling proportionally faster acquisition.^25^ The 60 kHz laser repetition rate delivered an on-sample fluence of ∼4 µJ/pulse, or ∼10 nJ/µm^2^, which was sufficient for bright fluorescence while preserving sample viability. As illustrated in Fig. 4, these imaging conditions enabled us to capture a broad portion of the root, while multiple cross-sectional planes (labeled as 1 - 4) extracted from different axial positions highlight clear cellular details within the epidermis, demonstrating subcellular axial resolution across the millimeter-scale volume. To confirm ST-LSM’s robustness, we repeated these measurements on fixed roots tips stained with propidium iodide (Fig. S4 including a brightfield image for anatomical reference), reaffirming the ST-LSM’s reliability across multiple sample conditions.

**Figure 4.**
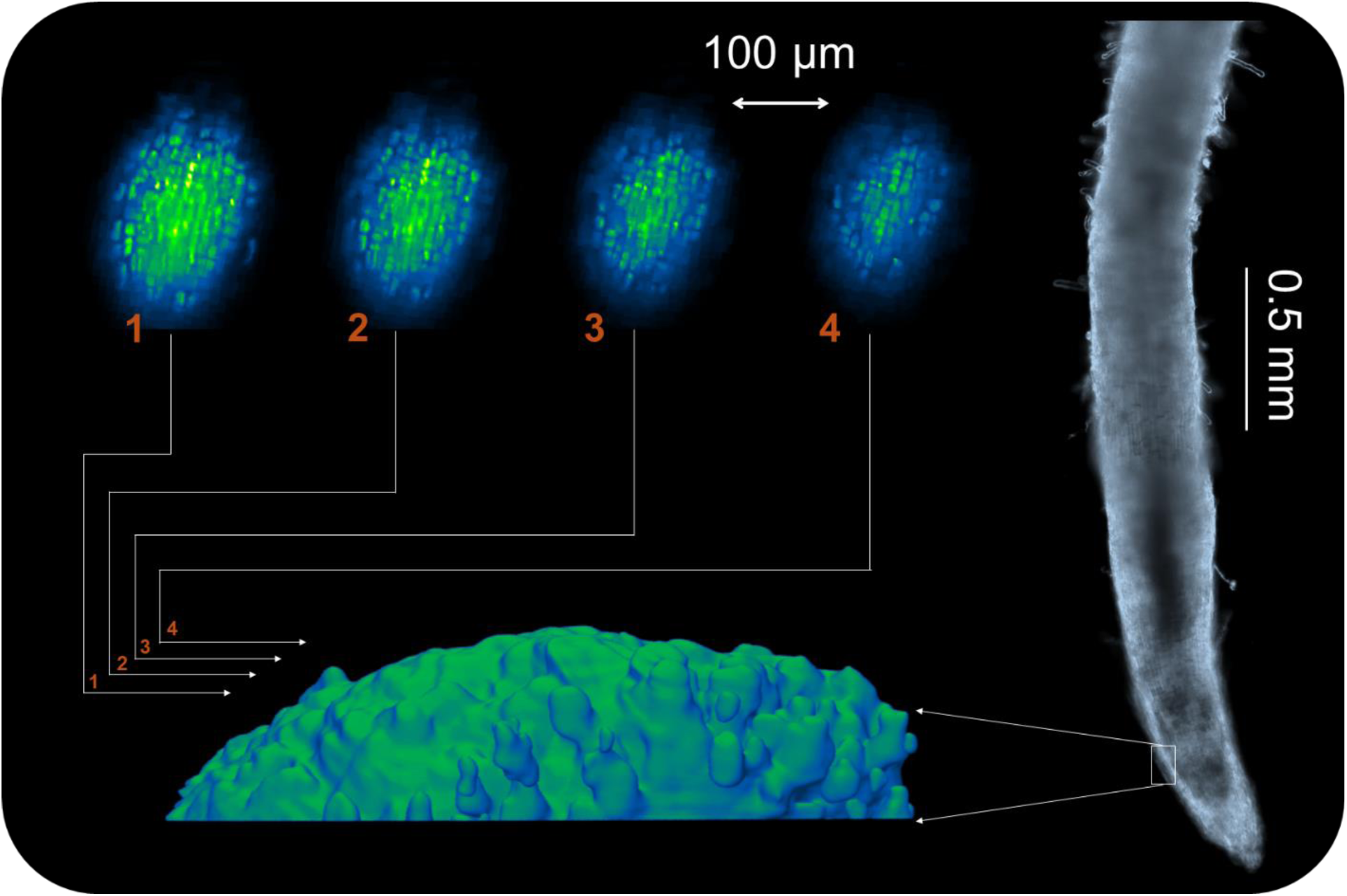
Root bioimaging enabled by ST-LSM. Volumetric reconstruction of a live *Medicago truncatula* root expressing a fluorescent protein. A widefield image (blue/gray) shows the entire root, with the boxed region is rendered in 3D using ST-LSM (green). Four representative cross-sections (1–4) illustrate internal structural detail over a depth of ∼400 μm.

Next, we applied ST-LSM to zebrafish embryos, a widely used vertebrate model in LSM/SPIM investigations, under similar fluence conditions as in the root experiments (∼10 nJ/µm^2^) at a 60 kHz repetition rate. As shown in Fig. 5a, we stained the fixed zebrafish embryos (prepared at 3.5 days post-fertilization, dpf) with propidium iodide to highlight cellular and nuclear distributions, enabling the clear identification of key anatomical features such as the yolk sac, spinal cord, swim bladder, and portions of the gastrointestinal tract. The extended propagation length of the STWP allowed us to capture large embryonic regions in a single view without refocusing, using x-axis stitching every 600 µm (Fig. 1c), where the total acquisition time scales with stitched field size rather than intrinsic beam formation constraints. Even delicate 3D structures remained intact and resolvable (Fig. 5a), reflecting the system’s capacity for comprehensive volumetric imaging. To highlight ST-LSM’s adaptability, we also imaged an embryo labeled with Nile Red (Fig. 5b), which revealed distinct lipid compartments not visible with propidium iodide, highlighting the broader utility and sensitivity of ST-LSM to varying fluorophores and staining proto­cols.

**Figure 5.**
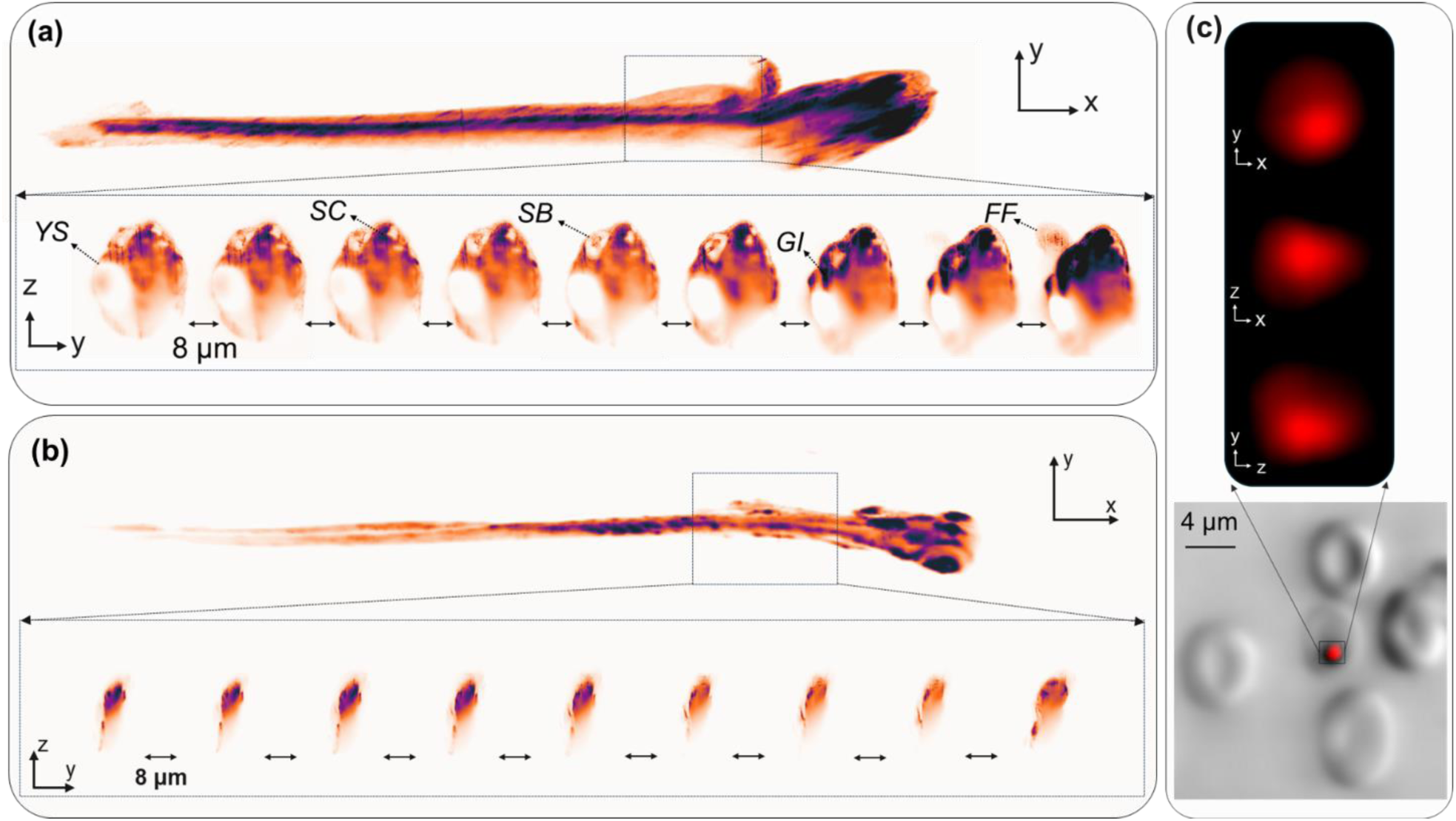
Seamless multiscale 3D bioimaging with a single ST-LSM setup from whole-organ structures to subcellular features. (a-b) Volumetric imaging of two *Danio rerio* (zebrafish) embryos stained with propidium iodide **(a)** and Nile Red **(b)**; *xy* views reveal gross morphology, while sequential *yz-plane* slices (8 μm spacing) highlight com­plementary structural details. In **(a)**, various organ structures are visible (*YS*: yolk sack; *SC*: spinal cord; *SB*: swim bladder; ***GI: gastrointestinal tract***; *FF*: fin fold, while in **(b)**, volumetric fluorescence imaging reveals the lipid distribution throughout the whole organ­ism. **(c)** A single red blood cell (RBC) infected by *Plasmodium falciparum*. Brightfield im­age (right) shows the RBCs; boxed region highlights an infected cell. Right: orthogonal fluorescence views (*xy*, *yz*, *xz*) of the parasite nucleus stained with propidium iodide, re­vealing its subcellular distribution.

Finally, we tested the subcellular resolving power of ST-LSM by examining human red blood cells (RBCs) infected with the malaria parasite P. falciparum, again employing similar fluence and repetition rates as in the root imaging case. As displayed in Fig. 5c, the ∼1.4 µm optical sectioning thickness provided strong axial confinement, enabling high-contrast 3D visualization of the parasite nucleus and clear differentiation of its intraerythrocytic localization. Collectively, these results highlight ST-LSM’s subcellular resolution, as well as how a single hardware setup can facilitate both whole-organism imaging, such as zebrafish embryos, and subcellular exploration (e.g., infected RBCs), broadening the scope of LSM/SPIM and offering powerful new opportunities in plant science, developmental biology, and pathogen research.

## Discussion

By harnessing the unique properties of STWPs, ST-LSM overcomes the long-standing constraints that have likely limited the broader adoption of LSM/SPIM, despite its exceptional resistance against phototoxicity and photobleaching. Specifically, we violated the innate sectioning resolution-FoV tradeoff, which forces the compromise between highly resolved but restricted imaging FoVs, or broader FOVs with lower resolution. By spatio-temporally structuring the illumination light-sheet, we eradicated this tradeoff, demonstrating light-sheets that maintain wavelength-scale thickness over millimeter-scale propagation distances. We note that this millimeter-scale propagation refers to the uniform illumination region of the light-sheet, whereas the instantaneous imaging FoV remains defined by the detection optics (objective NA and camera sensor size), as in conventional LSM. In this context, ST-LSM enables a 10× extension of the illumination FoV, allowing larger volumes to be imaged without compromising optical sectioning efficiency. To place this advancement in context, Table 1 provides a quantitative comparison of representative two-photon light-sheet microscopy modalities using literature-reported values. As detailed in Methods, pulse-duration measurements indicate a ∼40% increase under ST modulation, corresponding to an equivalent reduction in two-photon excitation efficiency at fixed pulse energy.

Crucially, under comparable two-photon illumination conditions, our ST-LSM configuration outperforms even the most advanced LSM/SPIM implementations that employ sophisticated beam-shaping strategies such as optical lattices, Bessel, and Airy light-sheets. First, ST-LSM delivers a further 2× increase in FoV under comparable two-photon illumination conditions.^37^ This stems from STWP light-sheets forming a class of optical fields that are effectively diffraction-free along one spatial dimension under experimentally accessible conditions.^29^ Bessel light-sheets still diffract, thus offering a smaller FoV than ST-LSM, while Airy light-sheets do not diffract but exhibit an asymmetric intensity profile that necessitates deconvolution.^6^ Fundamentally, monochromatic beam-shaping approaches, including lattice-based interference patterns, cannot achieve the diffraction-free lengths accessible to STWP light-sheets under two-photon excitation,^27^ even in presence of finite-energy constraints due to beam truncation.^5, 6, 10^ Indeed, the diffraction length of STWPs is determined by the spectral tilt angle θ along with the spectral uncertainty (the ‘thickness’ of the spatiotemporal spectrum; Fig. 2b) rather than the imaging aperture.^35^ Consequently, STWPs remain far less susceptible to aperture-induced shortening, preserving a robust propagation-invariance.

Second, Airy and Bessel beams typically carry strong sidelobes that intensify as the light-sheet is narrowed for sharper optical sectioning. ST-LSM introduces a tunable handle that balances main- and sidelobe intensities, thereby both expanding and streamlining the imaging parameter space. In this work, we explored a ∼0.4 µm range of light-sheet thicknesses selected to optimize two-photon imaging performance; this range does not represent a fundamental tuning limit. Third, Gaussian, Bessel, lattice, and Airy light-sheets are subject to Abbe’s sine condition, therefore requiring dual-objective configurations and high NA illumination lenses.^17^ These requirements, however, make them less adaptable to large or irregular specimens. In contrast, ST-LSM replaces the high-NA illumination objective with a simple cylindrical lens without sacrificing 3D spatial resolution. This enables a purely single-objective architecture that expands the illumination working distance by more than an order of magnitude relative to existing two-photon light-sheet implementations, as summarized in Table 1, without compromising spatial resolution . It also reduces system complexity while broadening compatibility with diverse samples, including microfluidics and specialized imaging chambers.

Taken together, these unique STWP attributes unlock multiscale investigations that bridge entire organisms, tissues, and single cells. By translating previously established STWP physics into an optimized SPIM-compatible optical-sectioning strategy implemented in a single-objective geometry, this work positions ST-LSM as a flexible, high-performance platform for diverse applications. We demonstrate this versatility here in ST-LSM imaging of plant roots, entire zebrafish embryos, and malaria parasite-infected RBCs. Finally, many other STWP attributes can be exploited in the context of LSM, including self-healing of STWPs after traversing scattering media,^27^ reduced speckle growth in scattering samples, and dispersion-free propagation in dispersive samples.^28^ These additional features raise the prospects of seamlessly folding additional modalities, including Raman and sec-ond- or third-harmonic microscopy, into the same setup.

By demonstrating that wavelength-scale axial resolution and millimeter-scale FoVs can coexist within a streamlined, purely single-objective design, we envision ST-LSM catalyzing and democratizing high-performance light-sheet imaging, accelerating investigations that have hitherto remained challenging due to the limitations of traditional modalities.

## Acknowledgments

We gratefully acknowledge funding from the U.S. Department of Energy, Office of Science, Office of Biological and Environmental Research (DE-SC0025418). We also acknowledge Maria J. Harrison (Boyce Thompson Institute) for preparing and providing the *Medicago truncatula* samples used in this research.

## Contributions

J.Z. performed all optical experiments. H.L. developed the theoretical model and numerical simulations. S.L. and D.M. prepared and supplied biological specimens. A.F.A. and M.K. optimized the STWP strategy and provided critical insight and data interpretation.

A.E.V. and D.N.C. supervised the project and drafted the manuscript. All authors discussed the results and approved the final version.

## Methods

### Microscopy setup

The experimental setup for generating ST wavepackets is shown in Fig. 2c, with the complete setup for synthesizing and utilizing these wavepackets in imaging is further detailed in Fig. S1. Briefly, we expanded the beam of a pulsed laser source (Carbide CB5-06 with i-OPA-TW-HP, Light Conversion center wavelength 1030 nm; repetition rate 60 kHz; 8 nm bandwidth, pulse duration ∼180 fs at the source) using two spherical lenses (focal lengths: 50 mm and 63 mm) arranged in a 4f configuration. This produced a spherically symmetric beam with a full width at half maximum (FWHM) diameter of ∼5 mm. We adjusted the beam intensity using a polarizer in combination with a half-wave plate (990-0060-11IR, EKSMA Optics). Following a beam splitter (CCM1-BS014, Thorlabs), the expanded beam was directed onto a reflective grating (33009FL01-530r, Newport Corporation) mounted onto a grating holder (DGA-25, Newport Corporation) placed at the focal distance (400 mm) of a cylindrical lens with curvature along the y-axis (Fig. 1a). The cylindrical lens directed the diffracted beam on to an SLM (E19×12-500-1200-HDM8, Meadowlark Optics) programmed with the characteristic 2D square phase profile of STWPs. Phase values ranging from 0 to 2π were encoded as 8-bit gray-scale levels from 0 to 255. Because the SLM operates in reflection, the modulated field retraced its path through the same cylindrical lens back to the diffraction grating, where the angularly dispersed spectrum was recombined into a single beam. The beam splitter then separated this retroreflected, spectrally encoded wave packet from the input path and directed it toward two orthogonal 4f relay systems composed of cylindrical lens pairs aligned along the y- and z-axes, respectively (Fig. S1). These two 4f systems relayed the phase pattern to the axial plane of an imaging objective. Specifically, the first 4f system used lenses with focal lengths f_y1_ = 1000 mm and f_y2_ = 500 mm, and the second used f_z1_ = 1500 mm and f_z2_ = 50 mm. To precisely incline the ST beam after these 4f relays, we incorporated a reflective prism (MIM-CUBE-III-KM, Applied Scientific Instrumentation) po­sitioned between lenses CL_y2_ and CL_z2_ (Fig. S1). The sample was mounted on a 3D linear stage (Applied Scientific Instrumentation) to allow precise alignment of the sample with the focal planes of the illumination lens and the imaging plane. For imaging, we deployed two water immersion objectives NIR APO 40×/0.8 (Nikon) and a 16.7×/0.4 (Special Op­tics) depending on the sample. Fluorescence images were acquired using a scientific CMOS camera (Zyla, Andor) positioned behind a short pass filter (ZFESH0750, Thorlabs). The camera has a physical pixel size of 6.5 µm. With the 40×/0.8 detection objective, this corresponds to an effective pixel size of 162.5 nm at the sample plane, and with the 16.7×/0.4 objective, 389 nm per pixel. No additional relay magnification was in­troduced in the detection path. Volumetric scanning was performed by scanning the mo­torized stages in the xz-plane (Fig. 1b) using a Tiger controller (Applied Scientific Instru­mentation) and Micromanager 1.4.^38^ Generally, the pulse energy at the sample plane was ∼10 nJ/μm^2^ (0.8 mW/μm^2^), with a total optical throughput from the laser source to the illumination lens of ∼10% (∼90% loss). The dominant loss (∼75%) arises from the non-polarizing beam splitter in the folded pulse-shaper geometry due to the double-pass configuration. Additional losses originate from the diffraction grating efficiency and the SLM modulation efficiency. These losses are not intrinsic to ST beam synthesis but are associated with the folded 4f pulse-shaper implementation used here. A linear (non-folded) shaper architecture would substantially improve transmission by eliminating the beam-splitter penalty. We adopted the folded configuration for alignment stability and because sufficient pulse energy was available for two-photon excitation under all reported imaging conditions. Pulse duration after the relay was measured using frequency-resolved optical gating (FROG; GRENOUILLE 8-50-USB, Swamp Optics). The Gaussian reference pulse (blank SLM) exhibited a duration of ∼212 ± 5 fs (mean ± SEM for n = 5 measurements), while the ST-modulated pulse measured ∼309 ± 10 fs under identical alignment conditions. This moderate increase is consistent with additional residual spectral phase when applying the ST mask, likely due to SLM imperfections (e.g., finite pixelation and phase quantization) that can degrade spectral recombination.

### Optical alignment

To ensure that the beam remained centered through all optical elements, we mounted cage systems along the optical path (Fig. S1). Once installed, the optical system remained stable for months, requiring only minor mirror adjustments every few days. To align the SLM pattern with the excitation beam, we positioned a CMOS camera (acA3800-14um, Basler) just before the pair of 4f systems. We further verified the alignment using the pulse shaper to generate a temporal Airy pulse,^34^ which we assessed with a frequency-resolved optical gating (FROG, GRENOUILLE 8-50-USB, Swamp Optics) measurement, as displayed in Fig. S6.

### Data acquisition and processing

A PC (Z8, Hewlett-Packard) equipped with Intel Xeon W-2123 W CPU 3.60 GHz processors and 128 GB RAM acquired and temporarily stored raw 3D images. Images were analyzed using Fiji on a workstation equipped with an Intel Core i7-7820X CPU @ 3.60 GHz processors and 128 GB RAM.^39^ 3D image representations were completed with the Avia (Leica) and Imaris (Andor) software.

### Sample preparation

To characterize the imaging FoV of ST-LSM, we employed both a Rhodamine 6G solution (∼0.1 mg/ml, AC419010050, Thermo Scientific Chemicals) and fluorescent YG particles (1 µm, 500 nm, and 200 nm diameter particles, 17154-10, 15700-10 and 15700-10,17151-10 Polysciences Inc.) embedded in an agarose gel. To image the particles, we used the NIR APO 40×/0.8 objective, while to characterize the light-sheet in the dye solution, we used the 16.7×/0.4 objective. To prepare the gel sample, we mixed 0.5% low temperature agarose (Ultra-Pure, Invitrogen) with water and kept the mixture in a convection oven at 80°C for 45 min until the agarose dissolved. Subsequently, we added the fluorescent YG microspheres of 1 μm diameter to the gel and mixed thoroughly. Prior to mixing, we diluted the particle solution of 1% solids for optimal particle density by approximately 10^4^. The mixture was poured into a custom holder between two coverslips for 15 minutes to solidify before imaging. We also imaged Medicago truncatula roots expressing an RFP-tagged membrane marker, using transgenic lines that constitutively express fluorescent reporters of subcellular compartments, as previously described.^40^ Root samples were prepared by scarifying M. truncatula seeds with sandpaper, sterilizing them in 30% Clorox with 0.1% Tween 20 for 5 minutes, rinsing three times with sterile distilled water, and placing them on sterile filter paper in a sealed Petri dish.^41^ Seeds were stratified at 4°C for 3 days, incubated at room temperature in the dark for 1 day, then transferred to light conditions (1,000 lx) for germination and growth. After 4 weeks, roots were mounted on glass coverslips and embedded in a thin layer of low melting point aga­rose (Thermo Fisher). The entire sample was immersed in water, with leaves kept above the water surface during imaging. In parallel, we imaged M. truncatula roots fixed in 50% ethanol and stained with 15 μg/ml propidium iodide (Invitrogen) to visualize cell walls. To image the roots, we used the 16.7×/0.4 (Special Optics) water-immersion objective. To prepare malaria parasite-infected red blood cells (RBCs), type O⁺ human blood (Grifols Bio Supplies, Los Angeles, CA, USA) was processed within one day of receipt. RBCs were isolated by mixing the blood with RPMI 1640 medium and centrifuging to remove white blood cells and plasma proteins. The washed RBCs were stored at 4°C for up to two weeks. Plasmodium falciparum (NF54 strain) cultures were initiated from frozen stocks, which were thawed at 37 °C and washed sequentially with decreasing concentra­tions of NaCl (12%, 1.6%, 0.9%) to remove glycerol. The resulting RBCs were transferred to flasks and cultured at 37°C in complete RPMI 1640 medium supplemented with 10% heat-inactivated human serum, HEPES, L-glutamine, hypoxanthine, and DL-lactic acid, using 4.0–6.0% hematocrit of washed type O⁺ RBCs. Cultures were maintained with daily medium changes and continuous gas exchange (5% CO₂, 5% O₂, 90% N₂). Parasitemia was monitored every two days by microscopic examination of Giemsa-stained thin blood smears. When parasitemia reached 10–12%, cultures were split and maintained under the same conditions. For imaging, infected RBCs were sampled from the cultures, fixed overnight in 2% paraformaldehyde at 4°C, washed three times with fresh RPMI 1640 me­dium, and stained with 100 μg/ml propidium iodide (Invitrogen) prior to imaging with the NIR APO 40×/0.8. To prepare the zebrafish (Danio rerio) embryo samples, adult fish were maintained on a 14:10 hour light:dark cycle in a recirculating, temperature-controlled system at 28°C at the University of Idaho under approved Institutional Animal Care and Use Committee (IACUC) protocols. Embryos were collected from breeder pairs at light onset, transferred to glass beakers with system water, and grown to approximately 3.5 days post-fertilization (dpf). For imaging, embryos were euthanized using MS-222 (tricaine) and fixed overnight at 4°C on a rocker in 4% paraformaldehyde prepared in phosphate-buffered saline (PBS). After fixation, embryos were washed once in PBS containing 1% Triton X-100, followed by three washes in PBS, and stored in PBS at 4°C until imaging. Prior to imaging, embryos were stained with 15 μg/ml propidium iodide (Invitrogen) or Nile Red (Thermo Fisher) in water and mounted on glass coverslips coated with low melting point agarose (Thermo Fisher) prior to imaging with the 16.7×/0.4 objective.

### Imaging FOV

To estimate the field of view (FoV), we imaged two-photon fluorescence excitation of a light sheet propagating through a diluted Rhodamine 6G solution (∼0.1 mg/mL). The FoV was quantified by extracting the fluorescence intensity profile along the x-axis (Fig. 2d) and fitting it with a Gaussian function in Origin Pro. The resulting fit parameters were used to compute the effective FoV that is further detailed in the man­uscript.

### Optical model

The phase mask design for ST wave packets. The relations between frequency and wavenumber are:

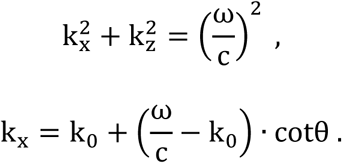

Combining the two above equations, one can obtain:

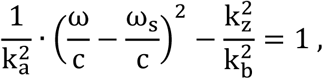

where 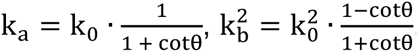, and 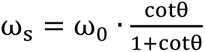.

For the phase mask in SLM, exp [iψ(y, z)], one can obtain:

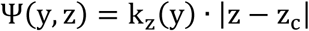

For each time frequency ω, it would be equipped with ±k_z_, which requires double-side phase modulation, i.e., |z − z_c_|.

Given that the dispersed profile z = z(λ) = z(ω), from the paraxial approximation, around the center ω_0_ (k_0_) one can find:

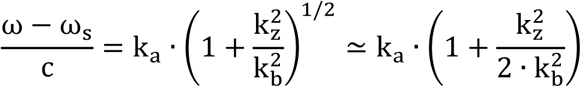

Simplify the above equation to:

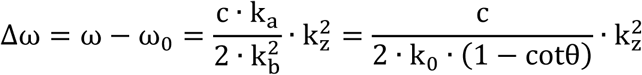

And combining with Δω = −2 · π · c · Δλ/λ_c_^2^, one acquires:

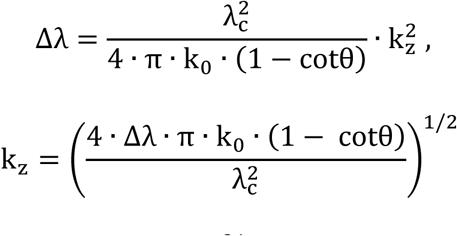

Compare the above relation to a scaling phase pattern:

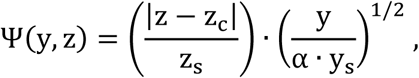

where z_s_ and y_s_ represent the scaling factors (actual full lengths on the SLM) along z and y directions, respectively. Then one can obtain the spectrum along y direction Δλ(y) = y/y_s_ · ΔΛ with ΔΛ being the whole wavelength range incident on the SLM, and in z axis for k_z_ modulation, the phase is given by:

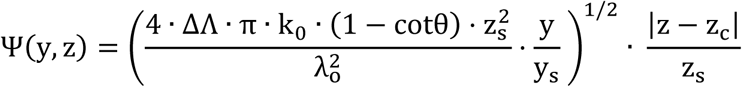

Given that, one can obtain 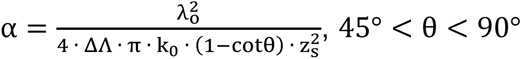, the curve of ω and k_z_ is hyperbolic. In this context, θ is the angle between the spatiotemporal spectral plane and the (k_x_) - axis in the (k_x_, ω/c) plane, determining the shape of the spectral trajectory on the light-cone that yields diffraction-free space-time light-sheets. Matlab was used to generate the STWP phase masks.^27^ Following the above methodology, we obtained the output 2D STWP profile and explored the relevant propagation properties given that the FWHM width of the input Gaussian beam is w_1_ = 5 mm and w_2_ = 5 mm with the wave­length λ_o_ = 1030 nm . The pertinent 4f systems provide de-magnifications of M_y_ = 1/2 or M_z_ = 1/30.

